# Targeting C42 in PKGIα alleviates diastolic dysfunction in HFpEF

**DOI:** 10.1101/2025.05.09.653217

**Authors:** Jie Su, Min Zhang, Xiaoping Yang, Mark Holt, Janice Raabe, Friederike Cuello, Ajay Shah, Michael J. Shattock, Joseph R. Burgoyne

## Abstract

**Background:** In our exploration of novel therapeutic strategies for treating heart failure with preserved ejection fraction (HFpEF), we focused on cysteine 42 (C42) in cGMP-dependent protein kinase Iα (PKGIα), which plays a crucial role in regulating blood pressure and diastolic relaxation. Our findings demonstrate that targeting C42 on PKGIα with urolithin A effectively limits diastolic dysfunction and cardiac remodeling in a multi-hit mouse model of HFpEF, while also improving the kinetics of relaxation and contraction in human-engineered heart tissue.

**Methods:** We evaluated various polyphenols, including urolithin A, for their capacity to oxidize and activate PKGIα. This assessment involved recombinant protein, isolated rat aortic smooth muscle cells, and myography experiments using isolated mesenteric vessels. Additionally, we investigated the ability of urolithin A to stimulate PKGIα-dependent phosphorylation of phospholamban (PLN) in neonatal rat ventricular myocytes (NRVMs). A multi-hit mouse model was employed to induce HFpEF in wild-type and C42S PKGIα knock-in (KI) mice. Following this, the mice were given vehicle or urolithin A by gavage, and heart function was evaluated using echocardiography. The relevance to human physiology was assessed using human-engineered heart tissue.

**Results:** The polyphenols quercetin, fisetin, and urolithin A were shown to induce C42-dependent activation of PKGIα. This mechanism of kinase activation led to vasorelaxation, which was attenuated in mesenteric vessels isolated from KI mice. Additionally, urolithin A treatment promoted PKGI-dependent phosphorylation of phospholamban in isolated cardiomyocytes. In a multi-hit model of HFpEF, urolithin A reversed diastolic dysfunction and cardiac remodeling in wild-type mice, but not in KI mice. These findings were shown to be likely relevant to humans, as urolithin A also enhanced the kinetics of relaxation and contraction in human-engineered heart tissue.

**Conclusions:** Targeting C42 in PKGIα with urolithin A presents a promising new strategy for the treatment of HFpEF.

## Introduction

Heart failure with preserved ejection fraction (HFpEF) accounts for nearly half of all heart failure cases and is associated with significant morbidity and mortality^1^. This prevalent condition is challenging to treat due to its complex pathophysiology, which is predominantly characterized by diastolic dysfunction. This dysfunction impairs ventricular relaxation and filling, often exacerbated during physical exertion, highlighting a limited cardiovascular reserve capacity, particularly in older individuals.

The cGMP-dependent protein kinase G Iα (PKGIα) plays a critical role in cardiovascular health via its activation by the canonical nitric oxide (NO)/cGMP pathway or through alternative mechanisms involving oxidants or 2’, 3’-cyclic GMP-AMP (cGAMP)^2, 3^. Oxidation of cysteine 42 (C42) on PKGIα by reactive oxygen species (ROS) or electrophilic compounds promotes disulfide dimerization, a process essential for regulating blood pressure and facilitating diastolic relaxation during myocardial stretch^4^. These important functions of C42 on PKGIα led us to hypothesize that this site could represent a novel therapeutic target to limit the progression of HFpEF.

Although natural polyphenols are often regarded as antioxidants due to their ROS-scavenging properties^5^, this view is oversimplified. Due to their redox cycling properties, polyphenols can paradoxically act as prooxidants by modifying protein cysteine thiols through Michael addition reactions^6^. This includes the formation of reactive quinones, which directly modify cysteine residues, and the generation of superoxide. Such modifications may underpin some beneficial effects of polyphenols. For instance, our previous research demonstrated that resveratrol mediates C42-dependent activation of PKGIα, which lowers blood pressure in hypertensive mice^3^.

In this study, we identified several polyphenols, including urolithin A, that potently induce C42-dependent activation of PKGIα. To evaluate the therapeutic potential of this mechanism, we utilized a multi-hit HFpEF model. This involved nephrectomy, implantation of a DOCA pellet, and a high-fat diet in mice. Urolithin A treatment significantly improved diastolic dysfunction and mitigated cardiac remodeling in this model, with both effects dependent on C42 activation of PKGIα. Furthermore, urolithin A enhanced relaxation and contraction kinetics in human-engineered heart (EHT) tissue, underscoring its translational potential.

## Methods

### Isolation and culture of rat aortic smooth muscle cells (VSMCs)

The thoracic aortas of rats were carefully isolated, with fat and connective tissue removed. The aortas were then incised and cut into small fragments, approximately 1-2 mm in size. These fragments were positioned lumen-side down in culture plates and submerged in a medium composed of Dulbecco’s Modified Eagle Medium (DMEM) (Gibco™, cat. #31966021) supplemented with 10% foetal bovine serum (FBS) (PAN-Biotech, cat. #P40-39500) and 1% penicillin/streptomycin (P/S) (Gibco™, cat. #15140122), to promote VSMCs migration onto the plate surface. After approximately five days, when VSMCs had proliferated and adhered to the plate, the aortic tissue pieces were discarded, and the cells were cultured in fresh medium until they reached confluence.

VSMCs were maintained at 37°C in a 5% CO_2_ humidified incubator and were regularly passaged. Rat aortic smooth muscle cells were passaged every 2-3 days when confluency reached >80% and cultured in DMEM media as above.

### Isolation and culture of neonatal rat ventricular myocytes (NRVMs)

Primary cultured cardiac ventricular myocytes were isolated from 1-2-day-old Sprague-Dawley rat pups (Charles River). Hearts were minced into ∼1 mm^3^ pieces in ice-cold ADS solution (6.8g/L NaCl, 4.76g/L HEPES, 0.12g/L NaH2PO4, 1g/L glucose, 0.4g/L KCl) and digested in 0.82g/L collagenase (Worthington™, cat. LS004176) and 0.6g/L pancreatin (SIGMA™, cat. #SLCD9444) at 37°C with agitation (200 rpm). Digestions were repeated 3-5 times, and supernatants were pooled in plating medium (68% DMEM media, Gibco™, cat. #31966021; 17% Medium 199, SIGMA™, cat. #M1264; 10% horse serum, 5% non-heat inactivated FBS, PAN-Biotech, cat. #P40-39500; 1% P/S, Gibco™, cat. #15140122; 2% 200mM glutamine, 1% non-essential amino acids).

Cells were filtered (100 µm), centrifuged (1150 rpm, 5 min), and pre-plated in T175 flasks for 2 hours to remove non-myocytes. The NRVM-rich supernatant was seeded onto gelatin (SIGMA™, cat. #G1393)-coated 24-well plates and incubated at 37°C. On Day 2, maintenance medium (DMEM/M199, 1% P/S, 2% glutamine) was replaced with plating medium (68% DMEM media, 17% Medium 199, 10% horse serum, 5% non-heat inactivated FBS, 1% P/S, Gibco™, 2% 200mM glutamine, 1% non-essential amino acids), and cells were cultured for 72 hours prior to treatments.

### Cell treatment

Media was changed to Earle’s Balanced Salt Solution (EBSS) (Gibco™, cat. #24010043) before cells were treated with 50 µM quercetin (Sigma-Aldrich, cat. #117-39-5), 50 µM fisetin (Universal Bio, cat. #CS-7840), 50 µM retinol (Sigma-Aldrich, cat. #R7632), 50 µM retinoic acid (Sigma-Aldrich, cat. #R2625), 50 µM urolithin A (Sigma-Aldrich, cat. #1143-70-0) or 50 µM ergothioneine (Sigma-Aldrich, cat. #497-30-3). Cells were supplemented with 1000x stocks of natural compounds, with vehicle also given at an equal volume to control wells.

To selectively inhibit PKGI activity, Cells were transfected with PKGI siRNA (Horizon Discovery, cat. #L-085799-02-0005) using Lipofectamine 3000 Reagent (Invitrogen, cat. #L3000008).

### Immunoblotting

Protein samples for electrophoresis were prepared in sample buffer (SB) (50 mM Tris-HCl pH 6.8, 2% w/v SDS, 10% glycerol, 0.0025% w/v bromophenol blue, with or without, 100 mM maleimide). SB was supplemented with 100 mM maleimide to make non-reducing samples, whereas for reducing samples 5% β-mercaptoethanol was added. 12 µL of each sample was loaded onto 15-well Mini-PROTEAN® TGX™ Gels (Bio-Rad, cat. #456-1086), then resolved at a constant 180V. Once the dye front had reached the bottom of the cassette, gels were removed from their cases. Using semi-dry transfer (Trans-Blot Turbo Transfer System, Bio-Rad), proteins were blotted onto polyvinylidene fluoride (PVDF) membranes (Bio-Rad). Subsequently, the PVDF membranes were blocked with 10 % non-fat dry milk or BSA in phosphate buffered saline (PBS) with 0.1 % v/v tween (PBS-T) for 1 hour, then incubated with the indicated primary antibody overnight at 4°C. After primary antibody incubation, membranes were washed with PBS-T 3 times for 10 min each, then incubated with the appropriate horseradish peroxidase (HRP)-linked secondary antibody for 1 hour at room temperature, followed by 3 washes with PBS-T as before. Samples were immunoblotted for Phospho-VASP (Ser239) (Cell Signaling, #3114), Phospho-PLN (Ser16)

(Badrilla, #A010-12AP), ANP (Proteintech, #27426-1-AP), a/b-Tubulin (Cell Signaling, #2148S) or GAPDH (Cell Signaling, #2118).

### PKGI activity assay

The activity of PKGI in response to urolithin A was measured using the PKGI Kinase Enzyme System (Promega, cat. #V4248) and the ADP-Glo™ Kinase Assay (Promega, cat. #V9101). PKGI was prepared following the manufacturer instructions, excluding the addition of dithiothreitol (DTT). 50 µM urolithin A was added to the corresponding wells of a 96-well plate. Reaction buffer containing the ATP/substrate mix was then added to each well, and the mixture was incubated for 60 minutes at room temperature. ADP-Glo™ Reagent was subsequently added to deplete residual ATP, followed by a 40-minute incubation at room temperature. Kinase Detection Reagent was next added to detect ADP, and after an additional 60-minute incubation, luminescence was measured using a GloMax® Discover Microplate Reader (Promega, cat. #GM3000).

### Myography

Mouse mesenteric vessels were dissected, isolated, and rings cut into ∼3 mm length in prewarmed Krebs buffer (126 mM NaCl, 2.5 mM KCl, 25 mM NaHCO3, 1.2 mM NaH2PO4, 1.2 mM MgCl2, 2.5 mM CaCl2, pH 7.2). The vessels were taken carefully to prevent endothelial damage due to stretch. The vessel rings were mounted with 25 µm wire in a Multi Wire Myograph System (DMT 620M) filled with Krebs buffer maintained at 37°C and bubbled with mixed gas (95% O_2_, 5% CO_2_). Optimal pre-tension was applied using the system’s normalization feature. After a 60-minute equilibration, the vessels were activated with 60 mM KCl to induce contraction, followed by three washes. After a 15-minute resting period, 10^−7^ M U46619 was used to induce contraction. In some experiments, vessels were pre-incubated with 5 µM ODQ for 3 hours. Following preincubation of U46619, a cumulative half-log dose-response curves for urolithin A (10^−7^ to 10^−4^ M) were applied to wild-type or C42 PKGI KI mesenteric vessels.

### Monobromobimane (mBBr) assay

Reactions with or without 200 µM urolithin A were made in the presence or absence of 50 µM cysteine in 100 mM Tris buffer at pH 8. After gently mixing, reactions were incubated at 37°C for 1 hour. After incubation 2 mM mBBr (ChemCruz™, cat. #sc-214629) was added to each sample and then incubated for a further 1 hour at 37°C. The reaction of mBBr with thiols results in a fluorescent adduct, which was measured at 394 nm excitation and 490 nm emission using a microplate reader (SpectraMax GeminiXS, Molecular Devices).

### Animals

All procedures were performed in accordance with the Home Office Guidance on the Operation of the Animals (Scientific Procedures) Act 1986 in the United Kingdom. Experiments were approved by the King’s College London Animal Welfare and Ethical Review Body. Mice used in this study were on the C57BL/6J genetic background. Male C42 PKGI KI mice or wild-type littermate controls were used in studies at 12-15 weeks of age. No animals were excluded from this study.

### Mouse model of HFpEF

The deoxycorticosterone acetate (DOCA)-salt hypertension model was used as it leads to diastolic dysfunction^7^. However, we employed a modification to this model where high-fat (HF) diet was included to induce a more robust and pathologically relevant model of HFpEF. The unilateral nephrectomy was performed, followed by subcutaneous implantation of a DOCA pellet (Innovative Research of America, cat. #M-121). On the day of DOCA-salt administration and unilateral nephrectomy, mice were provided water containing 1% NaCl and the standard diet was transitioned to a HF diet, concurrently with individual housing in cages equipped with running wheels. The Sham group underwent surgical procedures but without nephrectomy, DOCA implantation, NaCl slat water supplementation or high-fat diet. 21 days after initial generation of HFpEF, mice were administered urolithin A (50 mg/kg/d, dissolved in 0.25% CMC-Na) or vehicle control (0.25% CMC-Na) daily for 7 days by oral gavage. Mice of both genotypes were randomly subdivided into 2 groups per condition, resulting in establishment of 6 distinct experimental groups: 1) WT Sham; 2) WT HFpEF; 3) WT HFpEF + urolithin A; 4) C42S PKGI KI Sham; 5) C42S PKGI KI HFpEF; 6) C42S PKGI KI HFpEF + urolithin A. At the end of the protocol hearts were harvested and the blood in the chamber gently squeezed out with a small tweezer. The heart weight and the tibia length were then measured. The normalised heart weight (HW/TL) was calculated by dividing the heart weight by tibia length. Tissues were then flash frozen in liquid nitrogen for subsequent analysis.

### Echocardiography

Echocardiography was performed by high frequency linear array transducers (MX400, 30MHz, axial resolution: 50 μm, lateral resolution: 110 μm) equipped with a Vevo 3100 high resolution imaging system (FUJIFILM VisualSonics; Toronto, Ontario, Canada). Mice were anesthetised with 2% isoflurane to achieve constant and comparable heart rates (400-500 bpm) during image acquisition and imaged in the supine position on an electrical heating pad to maintain the core temperature around 37°C. Electrocardiographic (ECG) monitoring was carried out by limb electrodes. Subsequently, a standardized two-dimensional (2D) echocardiographic examination was generated to evaluate LV volumes and ejection fraction (EF), Pulsed wave Doppler flow and Tissue Doppler imaging were obtained in the apical 4-chamber view. All perspectives were archived as cine loops encompassing 300 frames.

The cardiac function was measured by echocardiography before DOCA administration, 21 days and 28 days after the implantation. All captured images were archived in raw DICOM format for subsequent offline analyses using the Vevo3100 LAB Software 5.7.1 (Visualsonics, Canada). Subsequent evaluations were conducted offline with blinding of mouse experimental groups.

### Histopathological analysis

The mouse heart was excised, and the ventricular tissues fixed overnight in 4% paraformaldehyde buffer. Samples were embedded in paraffin and cut into 5 µm thick slices. The slices were dewaxed and stained with wheat germ agglutinin (WGA) Alexa Fluor™ 488 conjugate (Invitrogen, cat. #W11261). Stained heart tissues were observed using a confocal microscope system (Leica TCS-SP5). The myocardial fibrotic area was assessed using Masson’s trichrome staining kit (Abcam, cat. #ab150686) following manufacturer instructions.

### Human-derived engineering heart tissue (EHT)

Cardiomyocytes were differentiated from the KOLF2 human induced pluripotent stem cell line using the StemMACS CardioDiff Kit (Miltenyi Biotec, cat. #130-125-28), according to the manufacturer’s instructions. Cells exhibiting wave-like contractions were dissociated from day 10 onward. Further experiments were performed at 14-25 days after initiation of differentiation.

Fibrin-based human-engineered heart tissue (EHT) were then generated as previously described^9^. Casting moulds were prepared in a 24-well plate format by using 2% agarose in PBS and a polytetrafluoroethylene (PTFE) spacer (EHT-Technologies, cat. #C0002), forming a mould for EHT embedding. After agarose solidification, transparent polydimethylsiloxane (PDMS) posts (EHT-Technologies, cat. #C0001) were inserted into the moulds. To form each EHT, 1 million cardiomyocytes were mixed with 5.6 µL 2x DMEM, 0.1% Y-27632 (10 mM; Sigma, cat. #Y0503-5MG), 2.5 µL fibrinogen (200 mg/ml in 0.9% NaCl; Sigma, cat. #F8630) and 97 µL NKM medium (DMEM, Sigma, cat. #D5796; 10% FBS, Sigma, cat. #F9665; 1% Penicillin/Streptomycin, Gibco, cat. #11548876; 1% L-Glutamine, Sigma, cat. #G7513) was prepared. Then, 3 µL thrombin (Sigma, cat. #T4648) was added to the master mix and quickly after mixing, it was pipetted into each agarose mould. The plate was incubated at 37°C with 7% CO_2_, 40% O_2_, and 98% relative humidity for 1.5-2 hours to facilitate fibrinogen polymerisation. After polymerisation, 500 µL of prewarmed NKM medium was added to overlay the EHTs for 30 minutes. EHTs were then carefully transferred to a new 24-well plate containing prewarmed EHT culture medium (DMEM, 10% horse serum, Sigma, cat. #H1138; 1% P/S; 10 µg/mL insulin, Sigma, cat. #I0516). Medium was changed three times per week by transferring the EHTs with their posts into 1.5 mL of fresh, prewarmed EHT medium.

Urolithin A (100 µM) or a vehicle control was added to engineered heart tissues (EHTs). Measurements were conducted during spontaneous activity at three time points: before treatment, and 30 minutes and 120 minutes after treatment. Contraction analysis of the EHTs was performed using the MuscleMotion software, focusing on contraction force and kinetics^10^. The contraction kinetics were assessed by measuring contraction and relaxation times at 90%, 50%, and 20% of peak tension (T_90%_, T_50%_, T_20%_).

The contractile force (F) was calculated based on the post-deflection degree (δ, µm), the silicone post length (L), radius (R), and the elastic modulus (E) of the PDMS. The calculation followed the formula^11^: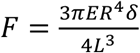.

### Statistical analysis

Data are shown as mean±SEM. Differences between groups were assessed using ANOVA followed by a two-tailed unpaired Student’s t-test or for multiple comparisons a Tukey’s test. Results were considered significant at a 5% significance level.

## Results

### Quercetin and fisetin mediate PKGIα C42 dimerization and enhance substrate phosphorylation

Natural compounds with potential thiol reactivity were evaluated for their ability to induce PKGIα disulfide dimerization and enhance kinase activity (Fig. 1A). Among the compounds tested, only quercetin effectively induced PKGIα disulfide dimerization. Consistent with this, quercetin also promoted substrate phosphorylation and synergized with cGMP to enhance kinase activation. Although retinoic acid, urolithin A, and ergothioneine did not induce PKGIα disulfide dimerization, they still enhanced substrate phosphorylation. Of these, only urolithin A synergized with cGMP, suggesting it may activate PKGIα through thiol oxidation.

**Figure 1.**
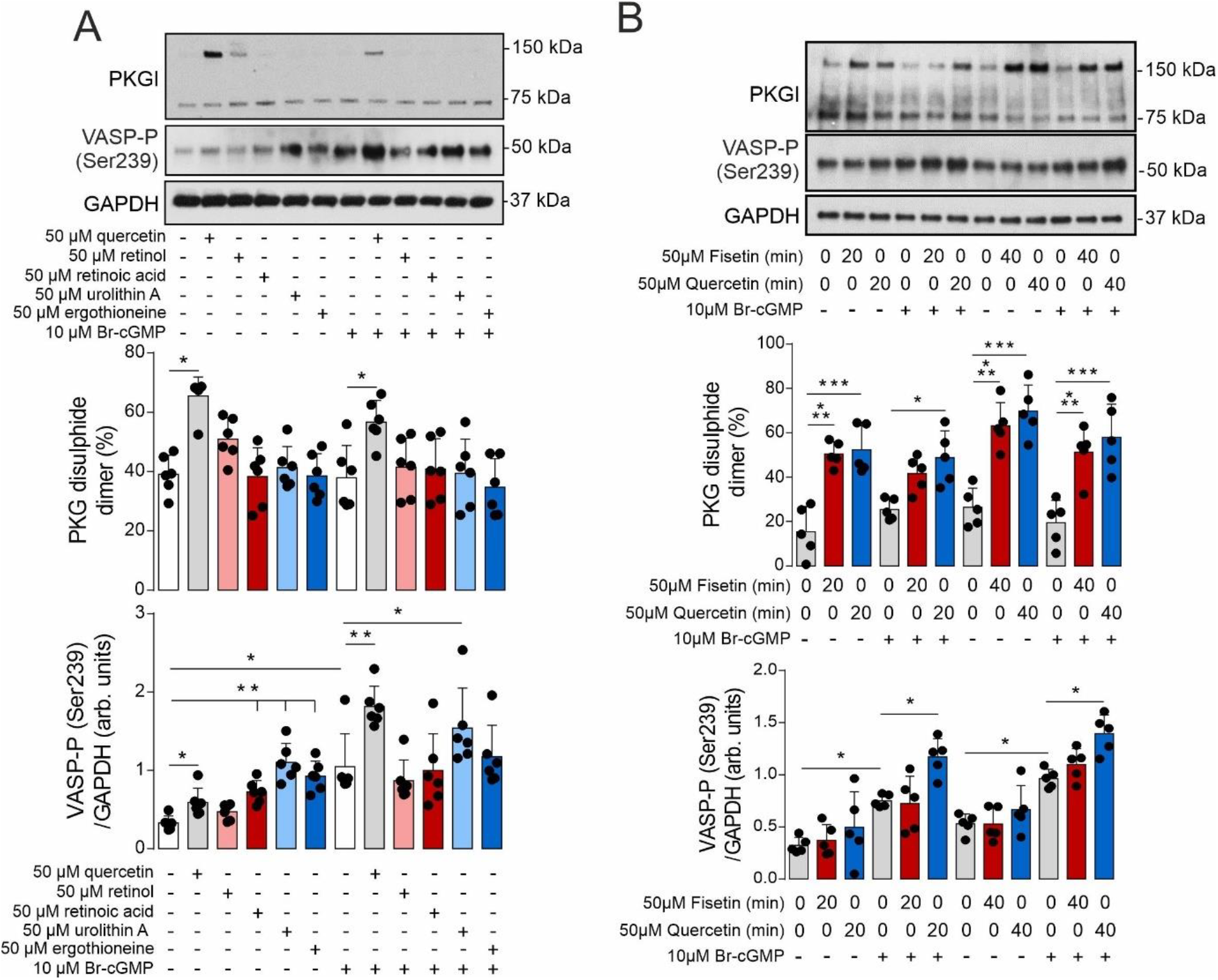
Fisetin and quercetin mediate PKGIα disulfide dimerization and increased substrate phosphorylation. **A**, PKGIα disulfide dimerization and VASP Ser239 phosphorylation in vascular smooth muscle cells treated with quercetin, retinol, retinoic acid, urolithin A or ergothioneine in the presence or absence of Br-cGMP (n=6). **B**, PKGIα disulfide dimerization and VASP Ser239 phosphorylation in vascular smooth muscle cells treated with quercetin in the presence or absence of Br-cGMP (n=5). *P<0.05; **P<0.01; ***P<0.005. Comparisons were made using 1-way ANOVA followed by the Tukey post hoc test.

Given the effectiveness of quercetin in inducing PKGIα disulfide dimerization, its activity was compared with the structurally similar flavonoid fisetin. Both compounds induced comparable PKGIα disulfide dimerization (Fig. 1B) and synergized with cGMP to enhance VASP phosphorylation, consistent with C42-dependent kinase activation.

### Urolithin A mediates C42-dependent PKGIα activation and vessel dilation

We next assessed the ability of quercetin and fisetin to modulate vasotone. Consistent with oxidative PKGIα activation, both compounds induced vasodilation (Fig. 2A). Quercetin was more effective in inducing vasodilation, and the mechanism of action was confirmed by comparing vessel responses from wild-type (WT) and PKGIα C42 knock-in (KI) mice. In vessels from KI mice, relaxation in response to quercetin was attenuated (Fig. 2B). To further validate the mechanism, soluble guanylate cyclase (sGC) was inhibited using ODQ. Inhibition of sGC did not affect relaxation to quercetin, supporting C42-dependent PKGIα activation (Fig. 2C).

**Figure 2.**
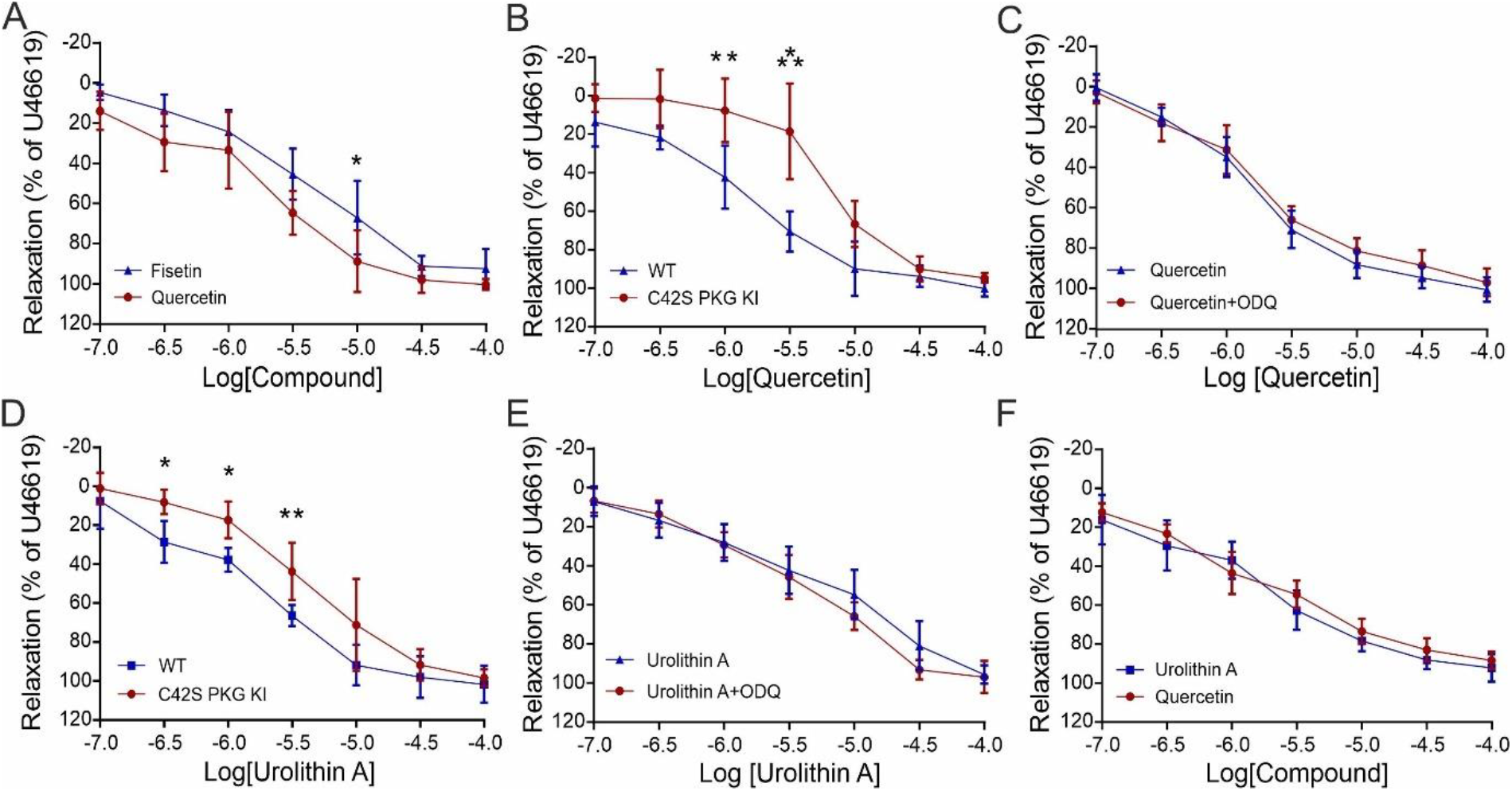
Quercetin and urolithin A induce PKGIα C42-dependent vessel relaxation. **A**, Comparing mesenteric vessel relaxation to quercetin or fisetin (n=5). **B**, Assessing relaxation of mesenteric vessels from WT and C42S PKG KI mice to quercetin (n=4). **C**, Assessing the impact of sGC inhibition on mesenteric vessel relaxation to quercetin (n=5). **D**, Comparing relaxation of mesenteric vessels from WT and C42S PKG KI mice to urolithin A (n=6). **E**, Assessing the impact of sGC inhibition on mesenteric vessel relaxation to urolithin A (n=5). **F**, Comparing mesenteric vessel relaxation to quercetin or urolithin A (n=4). *P<0.05; **P<0.01. Comparisons were made using 2-way ANOVA followed by the Tukey post hoc test.

Although urolithin A did not induce notable PKGIα disulfide dimerization, it still synergized with cGMP-dependent kinase activation, suggesting the potential for C42-dependent kinase activation. Therefore, we assessed its effect on vasotone and found that urolithin A mediated mesenteric vessel relaxation, which was also attenuated in vessels from KI mice (Fig. 2D). This suggests that, despite the lack of intermolecular disulfide formation, urolithin A likely activates PKGIα through oxidation of C42. This was further supported by experiments where sGC inhibition did not prevent relaxation by urolithin A (Fig. 2E). Given that both quercetin and urolithin A mediated PKGIα C42-dependent mesenteric vessel relaxation, we directly compared the two compounds. Both generated similar dose-dependent vessel relaxation (Fig. 2F).

### Urolithin A directly activates PKGIα which leads to enhanced cardiomyocyte phospholamban phosphorylation

Given the enhanced therapeutic potential of urolithin A compared to quercetin, including better bioavailability and existing human data^12-15^, we further investigated its mechanism of action. In *in vitro* kinase assays, urolithin A directly activated PKGIα, consistent with oxidation-dependent kinase stimulation (Fig. 3A). This finding was corroborated by assays measuring the thiol reactivity of urolithin A, which demonstrated its ability to compete with the alkylating agent monobromobimane (Fig. 3B). As urolithin A was likely to stimulate PKGIα through oxidation of C42, we hypothesized that this compound was modifying this residue to mimic the action of disulfide formation. This was substantiated in experiments where urolithin A limited the disulfide dimerization of PKGIα to diamide and led to detection of a urolithin A adduct on C42 (Fig. 3C, S1). Since C42 oxidation on PKGIα is linked to enhanced cardiac myocyte relaxation through phospholamban phosphorylation^4^, we next evaluated whether urolithin A could mediate this process. We found that urolithin A significantly increased cardiomyocyte phospholamban phosphorylation (Fig. 3D). Consistent with PKGIα activation, knockdown of this kinase attenuated phospholamban phosphorylation in cardiomyocytes treated with urolithin A (Fig. 3E).

**Figure 3.**
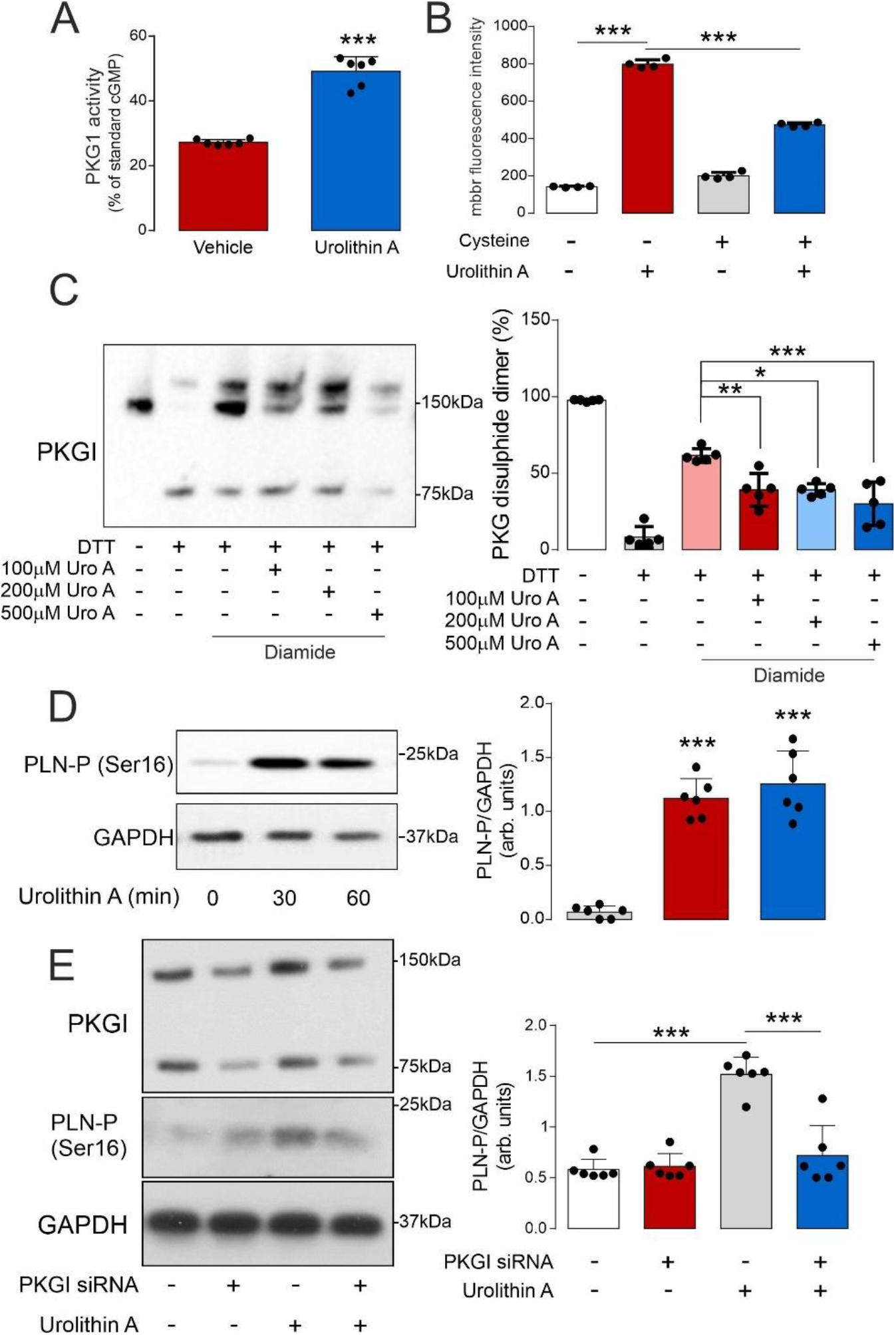
Urolithin A induces direct oxidation of PKGIα C42 to mediate kinase activation and enhance cardiomyocyte phospholamban phosphorylation. **A**, The activity of recombinant PKGIα in the presence or absence of urolithin A (n=6). **B**, Monobromobimane (mbbr) assay assessing the modification of cysteine by urolithin A (n=4). **C**, Treatment of reduced recombinant PKGIα with diamide in the presence or absence of urolithin A (n=5). **D**, Assessing PLN Ser16 phosphorylation in cardiomyocytes treated with or without urolithin A (n=6). **E**, Assessing the impact of knocking-down PKGI on urolithin A-dependent PLN Ser16 phosphorylation in cardiomyocytes (n=6). ***P<0.005. Comparisons were made using 1-way ANOVA (A, C, D) or 2-way ANOVA (B, E) followed by the Tukey post hoc test.

### PKGIα C42 oxidation by urolithin A limits diastolic dysfunction and cardiac remodeling

To evaluate the impact of urolithin A on myocardial dysfunction in HFpEF, we established a multi-hit preclinical model. In this model, mice underwent nephrectomy and were given 11-deoxycorticosterone acetate salt and a high-fat diet. To investigate the role of PKGIα C42, HFpEF was induced in both wild-type and KI mice (Fig. 4A-D, S2A-D). After HFpEF was established, mice were orally administered either vehicle or urolithin A for 7 days. In WT mice treated with urolithin A, left ventricular mass and heart weight were reduced (Fig. 4C, S2E) and diastolic function improved (Fig. 4D). These improvements in cardiac function and structure were accompanied by an increase in running distance (Fig. 4E, S2F), as well as a reduction in fibrosis and cardiomyocyte cell size (Fig. 4F, G, S2G). Notably, these effects were dependent on PKGIα C42, as they were not observed in KI mice. Thus, urolithin A limits cardiac dysfunction and remodeling in HFpEF through C42-dependent PKGIα activation.

**Figure 4.**
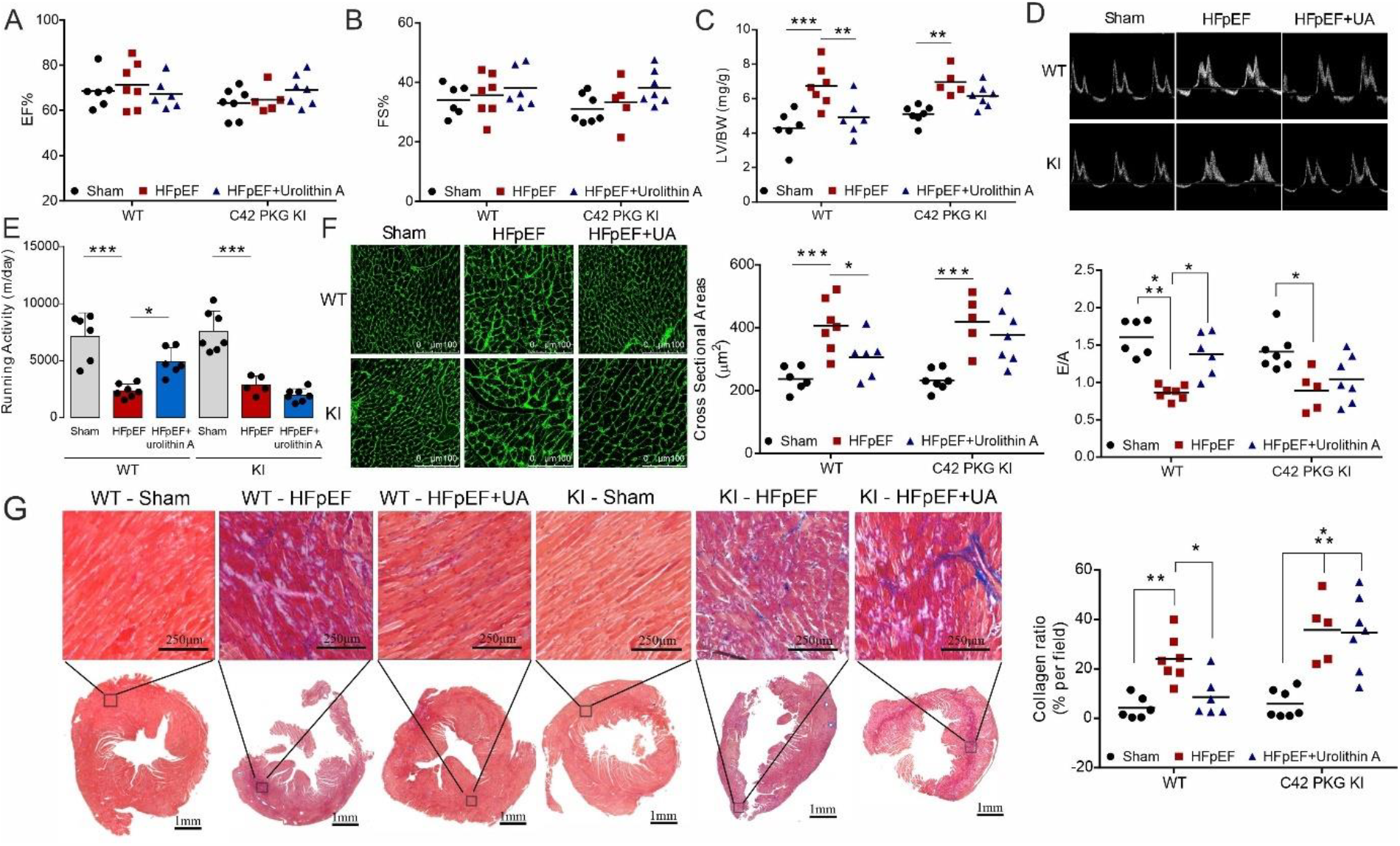
Urolithin A limits HFpEF that is dependent on PKGIα C42. Assessing **A**, EF, **B**, FS, **C**, LV/BW, **D**, E/A, **E**, running activity in WT and C42S PKGIα KI mice after sham surgery, HFpEF, of HFpEF with one week of diet supplementation with urolithin A (n=5-7). Assessing **F**, myocyte cross sectional area and **G**, collagen content in heart tissue excised from WT or C42S PKGIα KI mice after sham surgery, HFpEF, of HFpEF with one week of diet supplementation with urolithin A (n=5-7). *P<0.05; **P<0.01; ***P<0.005. Comparisons were made using 2-way ANOVA followed by the Tukey post hoc test.

### Urolithin A improves the kinetics of relaxation and contraction in human engineered heart tissue

To further evaluate the potential of urolithin A for treating HFpEF, we assessed its effects on contractile function in human engineered heart tissue. In tissues treated with urolithin A, there were no changes in maximal force generation or arrhythmias detected (Fig. 5A). However, there was a significant improvement in the kinetics of contraction and relaxation (Fig. 5B, C). Together, these findings suggest that urolithin A can enhance contraction and relaxation parameters in human cardiac tissue.

**Figure 5.**
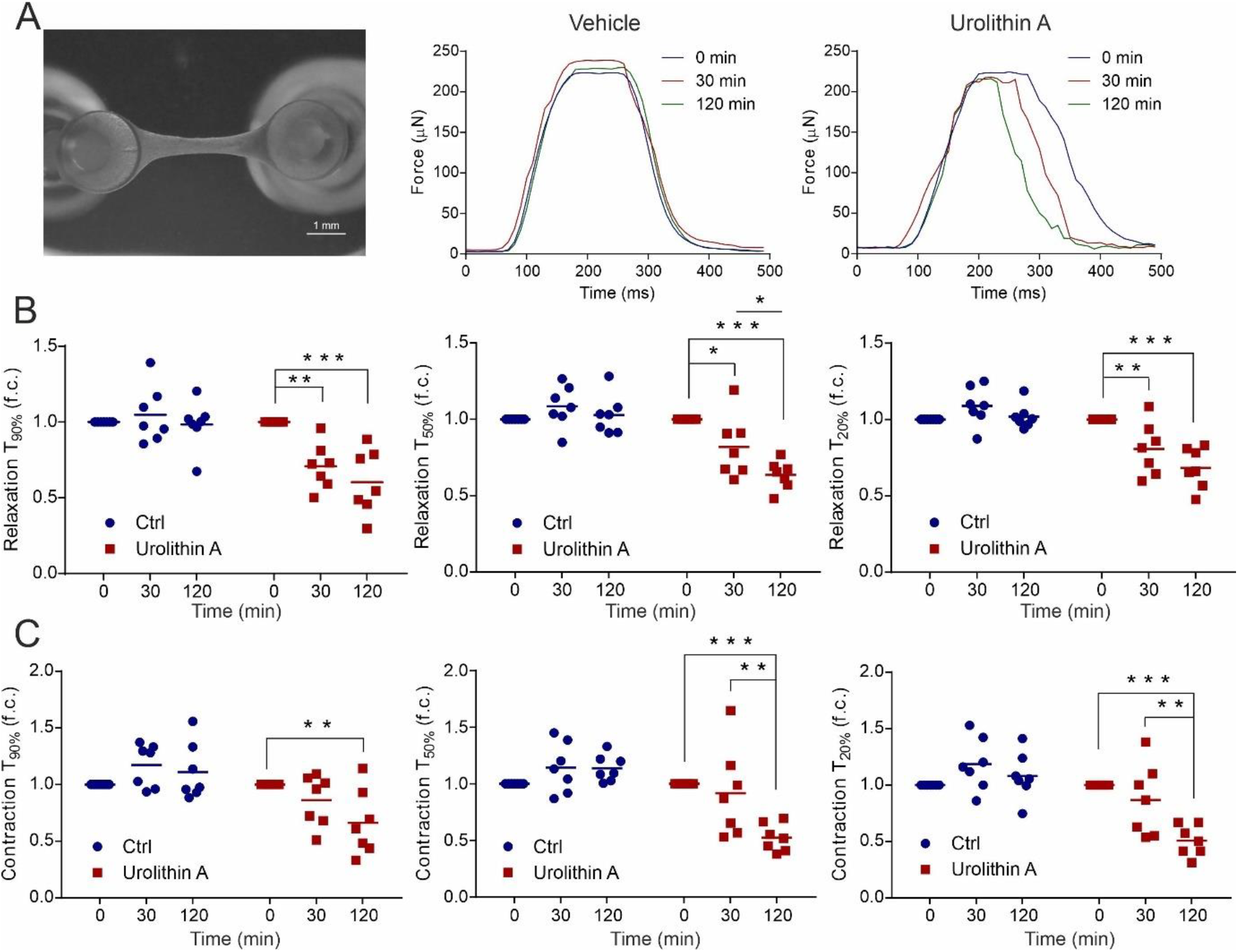
Urolithin A improves relaxation and contraction kinetics in engineered human heart tissue. **A**, Representative force generation in EHTs treated with vehicle or urolithin A. **B**, Kinetics of relaxation in EHTs treated with or without urolithin A (n=6). **C**, Kinetics of contraction in EHTs treated with vehicle or urolithin A (n=6). *P<0.05; **P<0.01; ***P<0.005. Comparisons were made using 2-way ANOVA followed by the Tukey post hoc test.

## Discussion

Impaired nitric oxide bioavailability and reduced PKGI activity are associated with the pathology of HFpEF, as supported by clinical evidence^16-18^. Based on this understanding, therapeutic strategies targeting this pathway have been explored using the phosphodiesterase type 5 (PDE5) inhibitor sildenafil and soluble guanylyl cyclase (sGC) stimulator vericiguat^19, 20^. However, these approaches failed to improve the primary outcomes or secondary endpoints in clinical trials. Several factors may explain these outcomes. Endothelial dysfunction in HFpEF patients likely results in reduced NO and natriuretic peptide bioavailability^21, 22^, diminishing the efficacy of PDE5 inhibition. Additionally, the VITALITY trial using vericiguat may have included a less symptomatic HFpEF population, which could have influenced the results. Alternatively, it is possible that NO signaling is not central to the progression of HFpEF, a hypothesis supported by the failure of oral nitrates and inhaled nitrites to improve clinical outcomes in this population^23, 24^. Our previous work demonstrated that cGMP mitigates C42-dependent PKGI activation^25^, a process known to enhance diastolic relaxation^4^. This suggests that the limited efficacy of NO-targeted therapies may arise from mechanisms beyond impaired NO signaling. Consequently, we hypothesized that directly targeting C42-dependent activation of PKGI could alleviate HFpEF progression through a dual mechanism by mediating vasodilation while directly improving diastolic function. This rationale led to the identification of a novel therapeutic strategy for HFpEF by targeting C42 in PKGIα using urolithin A.

Several natural compounds were selected for their potential to activate PKGIα via C42 thiol oxidation. These compounds were chosen based on their intrinsic α,β-unsaturated carbonyl groups or their ability to form such modifications through redox cycling. The presence of an α,β-unsaturated carbonyl group provides a capacity to modify cysteine thiols through a Michael addition reaction^26^. Among the selected compounds, quercetin and its structurally similar flavonol, fisetin, were particularly effective in inducing PKGIα disulfide dimerization and enhancing kinase activation. Additionally, urolithin, which did not promote intermolecular C42 disulfide formation, significantly enhanced PKGIα activation and demonstrated synergy with cGMP. This finding highlighted the potential for urolithin A to also mediate redox-dependent kinase activation. This would align with our previous findings, where the C42-dependent PKGIα activator G1 also synergized with cGMP^27^. The likely mechanism for this synergy involves disulfide formation, which enhances substrate interaction, while cGMP directly activates the kinase.

Consistent with the synergistic activation of PKGIα both quercetin and urolithin A were able to induce vasodilation that was dependent on C42. As urolithin A did not induce notable detection of the C42 PKGIα intermolecular disulfide this indicated a different mechanism of kinase activation. It was rationalized that this was likely through formation of a urolithin A adduct on C42. Consistent with the adduction of urolithin A on C42 this compound was found to be thiol reactive and could induce direct PKGIα activation. Furthermore, urolithin A limited the diulfide dimeirzation of PKGIα mediated by diamide, thus consistent with a urolithin A adduct on C42, which was confirmed by mass spectrometry analysis. It is likely that urolithin A is able to adduct to C42 through redox cycling and formation of a thiol reactive quinone intermediate.

Given its bioavailability and the health benefits associated with oral consumption of urolithin A in humans, this compound was further explored as a potential therapy for HFpEF^12-14^. In humans, urolithin A has a favorable safety profile, and its bioavailability has been shown to improve mitochondrial and cellular health in elderly individuals^13^. Additionally, a study found that urolithin A supplementation was safe, well-tolerated, and that long-term exposure likely counteracts age-associated muscle decline by improving muscle endurance and plasma biomarkers^14^. In mice, oral administration of urolithin A in obese mice alleviated metabolic cardiomyopathy by activating mitophagy^28^.

In cardiomyocytes, urolithin A induced PKGIα-dependent phosphorylation of PLN. This is consistent with the preferential phosphorylation of PLN by the C42-oxidized form of PKGIα^29^. Phosphorylation of PLN at Ser16 relieves its inhibitory effect on SERCA2a, enhancing the uptake of Ca^2+^ into the sarcoplasmic reticulum^30^. This mechanism of Ca^2+^ regulation by PKGIα contributes to diastolic relaxation and is involved in myocardial stretch, where it fine-tunes the Frank–Starling response^4^. Consistent with the ability of urolithin A to modulate PKGIα through C42 oxidation, this compound was found to limit the severity of HFpEF. Additionally, using redox-dead C42S knock-in (KI) PKGIα mice, we confirmed that the beneficial effects of urolithin A were primarily mediated through the oxidation of this residue. It is likely C42 thiol-dependent activation of PKGIα provides a dual benefit as it will not only mediate vasodilation but also directly enhance diastolic function. Together, these mechanisms limit cardiac remodeling and dysfunction.

To evaluate the translational potential of urolithin A in humans, we assessed its impact on the functional parameters of engineered human heart tissue. Urolithin A was found to improve the kinetics of both relaxation and contraction. Together with these findings, our results identify urolithin A as a novel thiol-dependent activator of PKGIα, which limits the severity of HFpEF in a preclinical model and is likely to improve cardiac function in patients with this condition.

